# Federated Knowledge Retrieval Elevates Large Language Model Performance on Biomedical Benchmarks

**DOI:** 10.1101/2025.08.01.668022

**Authors:** Janet Joy, Andrew I. Su

**Author notes:** Correspondence: Janet Joy and Andrew I. Su.

## Abstract

**Background:** Large language models (LLMs) have significantly advanced natural language processing in biomedical research, however, their reliance on implicit, statistical representations often results in factual inaccuracies or hallucinations, posing significant concerns in high-stakes biomedical contexts.

**Results:** To overcome these limitations, we developed BTE-RAG, a retrieval-augmented generation framework that integrates the reasoning capabilities of advanced language models with explicit mechanistic evidence sourced from BioThings Explorer, an API federation of more than sixty authoritative biomedical knowledge sources. We systematically evaluated BTE-RAG in comparison to traditional LLM-only methods across three benchmark datasets that we created from DrugMechDB. These datasets specifically targeted gene-centric mechanisms (798 questions), metabolite effects (201 questions), and drug–biological process relationships (842 questions). On the gene-centric task, BTE-RAG increased accuracy from 51% to 75.8% for GPT-4o mini and from 69.8% to 78.6% for GPT-4o. In metabolite-focused questions, the proportion of responses with cosine similarity scores of at least 0.90 rose by 82% for GPT-4o mini and 77% for GPT-4o. While overall accuracy was consistent in the drug–biological process benchmark, the retrieval method enhanced response concordance, producing a greater than 10% increase in high-agreement answers (from 129 to 144) using GPT-4o.

**Conclusion:** Federated knowledge retrieval provides transparent improvements in accuracy for large language models, establishing BTE-RAG as a valuable and practical tool for mechanistic exploration and translational biomedical research.

## 1 Introduction

Large language models (LLMs) have rapidly advanced the state of natural-language processing, reaching or surpassing expert performance across a wide range of biomedical tasks, including cell type annotation, protein-structure prediction and automated synthesis of clinical-trial results ^1–6^. However, the underlying generative methodology of these models, which sequentially predict tokens based on statistical patterns learned from massive text corpora, renders them susceptible to hallucinations, defined as outputs that are syntactically fluent yet factually incorrect ^7,8^. Such inaccuracies pose significant risks in biomedicine, where even minor errors can misdirect research efforts, delay critical therapeutic discoveries, or compromise patient safety ^7,9–11^. Indeed, recent assessments underscore that hallucination rates remain too high for safe and effective deployment in clinical and research-intensive environments^12,13^.

Efforts to mitigate these hallucinations through domain-specific pre-training and prompt engineering have yielded only incremental improvements, as these approaches continue to embed knowledge implicitly within opaque model parameters and fail to reliably surface evidence provenance ^14–16^. Retrieval-augmented generation (RAG) has emerged as a promising solution, explicitly grounding model-generated responses by dynamically incorporating external, verifiable evidence into prompts ^17–19^. Within biomedical question-answering contexts, RAG approaches consistently reduce hallucinations and elevate factual accuracy compared to parameter-only models. Nonetheless, the efficacy of RAG hinges critically on the precision, comprehensiveness, and currency of the retrieved contextual evidence ^20–22^.

Knowledge graphs (KGs) are particularly compelling resources for RAG because they explicitly represent biological entities and their relationships, support multi-hop mechanistic reasoning, and maintain persistent identifiers that simplify provenance tracking ^23–26^. Yet most biomedical KGs are tuned to a narrow slice of biology (for example, protein–protein interactions) or require extensive curation to remain current, limiting their utility for cross-domain mechanistic reasoning. To address these challenges, BioThings Explorer (BTE) integrates and federates 61 authoritative biomedical APIs into a continuously updated meta-knowledge graph that encompasses genes, pathways, drugs, diseases, phenotypes, and more ^27^. The API-centric framework of BTE returns structured JSON triples annotated with semantic types and evidence citations from reputable biomedical databases such as Gene Ontology, DrugBank, and Pubmed central using Translator Reasoner API (TRAPI) specification ^28–30^.

Here, we introduce BTE–RAG (BioThings Explorer–Retrieval-Augmented Generation), a novel framework that integrates the conversational fluency and reasoning capabilities of advanced LLMs with the explicit, multi-domain mechanistic knowledge captured by BTE. BTE–RAG dynamically executes targeted, query-focused graph traversals to retrieve concise, mechanistically pertinent evidence, formulates this evidence into declarative context statements, and augments model prompts accordingly.

To rigorously assess the performance of BTE-RAG in biomedical question answering, we systematically created three specialized benchmark datasets from DrugMechDB, a curated knowledge base containing 5,666 expert-annotated mechanistic pathways with literature validation ^31^. These datasets consist of gene-centric (n = 798), metabolite-centric (n = 201), and drug-centric (n = 842) question–answer pairs, each explicitly reflecting the causal flow from drug through intermediate biological nodes to disease outcomes. Across all three DrugMechDB-derived benchmarks, BTE–RAG robustly improves factual grounding, accelerates convergence to correct responses over diverse biomedical entities relative to an LLM-only baseline.

Collectively, these findings establish BTE–RAG as a powerful, practical tool for reducing hallucination risks and enhancing mechanistic clarity, significantly advancing the transparency, reliability, and utility of language model-driven biomedical discovery and clinical decision-making.

## 2 Materials & Methods

### 2.1 BTE–RAG Framework and Baseline Comparison

The BTE–RAG framework systematically compares two distinct inference routes to evaluate the impact of structured, mechanistic context on large language model (LLM) outputs (Figure 1A). The first inference route, labeled “LLM-only,” directly submits user-generated questions to the language model without external context augmentation. The second route, labeled “BTE–RAG,” integrates structured mechanistic evidence retrieved from BioThings Explorer prior to submitting an enriched, evidence-supported prompt to the same language model. This dual-path design allows rigorous evaluation of how explicitly retrieved context influences both answer accuracy and the factual grounding of model-generated responses. The BTE–RAG architecture comprises three key phases: entity recognition, knowledge-graph-based retrieval via BTE, and generative inference utilizing context-augmented LLM prompting.

**Figure 1:**
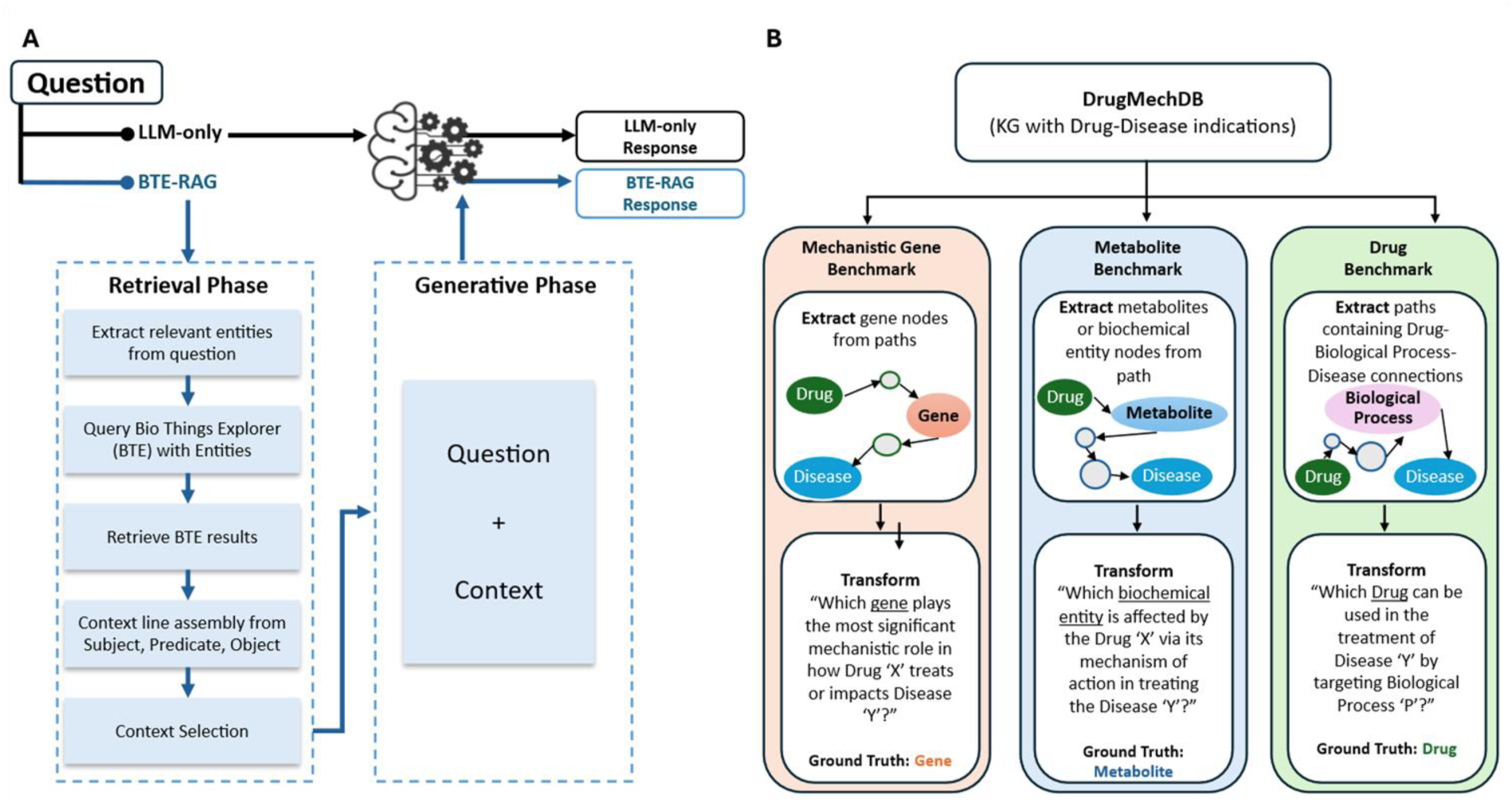
Retrieval–Augmented Generation workflow and derivation of mechanistic evaluation benchmarks. **(A)** Schematic of the BTE-RAG pipeline, which augments large language model (LLM) responses with context retrieved from the BioThings Explorer (BTE) knowledge graph. In the LLM-only pathway, the model generates a response using only the input question. In contrast, BTE-RAG operates in two phases: a Retrieval Phase, where relevant entities are extracted from the question and queried against BTE to collect mechanistically relevant subject–predicate–object triples, and a Generative Phase, where this curated context is appended to the input question and passed to the LLM. The resulting outputs: LLM-only or BTE-RAG, can be directly compared to assess the impact of knowledge-augmented generation. **(B)** Construction of benchmark datasets from DrugMechDB, a curated biomedical knowledge graph of drug–disease mechanisms. Directed paths connecting a drug to a disease were mined and transformed into structured questions targeting different mechanistic facets: (i) gene nodes (Mechanistic Gene Benchmark), (ii) biochemical entities or metabolites (Metabolite Benchmark), and (iii) drug–biological process– disease paths (Drug Benchmark). Each benchmark provides paired questions and gold–standard labels for rigorous, domain–specific evaluation of retrieval–augmented generation.

#### Entity Recognition

The retrieval phase begins with precise identification of biomedical entities mentioned within each input question. For the current benchmarks, entities such as drugs, diseases, metabolites, and biological processes were pre-annotated and standardized to established knowledge graph identifiers, enabling automated recognition at runtime. Additionally, the framework includes a zero-shot entity extraction module that leverages a specialized task-oriented prompting approach. This module is currently optimized for retrieving drugs and diseases from queries, with potential to extend extraction capabilities to include other biomedical entities as needed.

#### Knowledge Graph Retrieval

Identified biomedical entities are translated into structured queries interfacing directly with BTE. BTE integrates 61 authoritative biomedical databases under a unified knowledge graph schema, accessible via the programmatic API endpoint (/v1/query). Each query to BTE specifies an input entity (e.g., disease, drug, or biological process) along with desired output entity categories, following the TRAPI query format. In response, BTE returns structured JSON data that includes a detailed knowledge graph containing two key components: “nodes,” which describe biomedical entities along with their semantic categories and standardized names; and “edges,” which specify the explicit relationships (predicates) between pairs of entities, supplemented by provenance details indicating the primary knowledge sources.

For each benchmark dataset, targeted queries were structured to retrieve mechanistically relevant context. Specifically, in the gene-centric benchmark, queries separately utilized disease and drug entities to retrieve directly linked gene and protein nodes. In the metabolite-centric benchmark, disease and chemical (drug) entities were queried independently to identify connected biochemical entities. For the drug– biological process benchmark, separate queries using disease entities and biological process entities were conducted to retrieve associated chemical entities (drugs). Upon receiving the structured knowledge graph responses from BTE, both node and edge information were systematically processed. Nodes were extracted along with their semantic categories and descriptive names, while edges were parsed to identify subject-object pairs, predicates, and associated primary knowledge sources. Nodes and edges were subsequently merged to construct coherent statements that succinctly describe each mechanistic relationship (e.g., “drug X inhibits gene Y”). These concise, natural-language context statements collectively formed the mechanistic evidence provided to the language models during the generative inference phase, significantly enhancing the transparency, interpretability, and accuracy of the generated outputs. Supplementary Figure S1 provides a detailed schematic illustrating the complete BTE– RAG pipeline workflow, demonstrating a representative query and the subsequent processing and integration steps.

#### Context Selection

Two distinct evidence-inclusion strategies were systematically assessed for each question. The first strategy incorporates the entire set of sentences retrieved by BTE, leveraging the extensive 128,000-token context window of GPT-4o ^32^. The second strategy employs sentence-level cosine similarity filtering using ‘S-PubMedBert-MS-MARCO’ embeddings, retaining only sentences whose similarity scores with the query exceed a predefined percentile threshold ^33^. Running these two strategies concurrently enables a direct evaluation of the impact of comprehensive versus selectively pruned contextual evidence under identical experimental conditions.

#### Generative Inference

For the generative phase, selected context sentences and the original query were concatenated to form an enriched prompt submitted to both GPT-4o and GPT-4o-mini models. Models were configured deterministically (temperature set to 0) to produce reproducible outputs. Parallel runs of the LLM-only baseline used identical questions without the BTE-derived context. To streamline downstream analyses and ensure objective comparisons, language models were instructed explicitly to output structured JSON responses devoid of extraneous explanatory text. Detailed system prompts are shown in Supplementary Table S1.

#### Evaluation

Generated outputs were benchmarked against curated ground-truth annotations from the constructed mechanistic datasets. For entity-specific tasks (e.g., gene identification), correctness was evaluated via exact, case-insensitive string matching. For semantically nuanced responses (e.g., metabolites and drugs), BioBERT-based embeddings (‘BioBERT-mnli-snli-scinli-scitail-mednli-stsb’) quantified the semantic similarity between model-generated outputs and reference answers ^34^. Answers surpassing a predetermined similarity threshold were classified as accurate. Collectively, these standardized evaluation methodologies ensure scalable, objective, and reproducible assessment of the fidelity and biological coherence of model predictions, rigorously testing the utility and impact of knowledge graph-enhanced prompting in biomedical reasoning contexts.

### 2.2 Datasets from DMDB

### 2.2 Construction of Mechanistic Question–Answer Benchmarks from DrugMechDB

DrugMechDB is a rigorously curated biomedical knowledge graph designed to represent therapeutic mechanisms through explicit stepwise paths. These pathways originate from drug nodes, traverse biologically meaningful intermediate entities, and culminate at disease nodes, collectively delineating mechanisms underlying drug–disease interactions ^31^. The current version of DrugMechDB contains 5,666 curated mechanistic pathways, providing comprehensive coverage for 4,583 distinct drug–disease indications. Each node within DrugMechDB is systematically mapped to a standardized Biolink category and anchored to stable identifiers, while each relationship (edge) is annotated with a controlled predicate ^35^. This structured, granular, and provenance-rich resource enables robust benchmarking of computational models focused on mechanistic inference rather than simple associative or co-occurrence patterns.

To comprehensively evaluate the BTE–RAG framework across multiple levels of biological resolution, DrugMechDB was systematically transformed into three complementary mechanistic question–answer (QA) benchmarks, each highlighting a distinct biological focus: genes, metabolites, and drugs (Figure 1B).

Gene-Centric Benchmark: Mechanistic pathways were initially filtered to retain those containing exactly one internal node annotated as a Gene entity. Gene identifiers were resolved into standardized HGNC symbols using MyGene.info services; pathways containing deprecated or ambiguous identifiers were systematically excluded ^36^. Each remaining mechanistic pathway was converted into a structured question of the form: “Which gene plays the most significant mechanistic role in how Drug ’X’ treats or impacts Disease ’Y’?” The corresponding HGNC gene symbol served as the definitive ground truth. Following deduplication across different indications, this dataset comprised 798 unique QA pairs.

Metabolite-Centric Benchmark: To capture downstream biochemical effects, pathways exclusively containing taxonomic relationships (such as “subclass” predicates) were removed to ensure mechanistic specificity. Selected pathways included exactly one metabolite node, identified specifically by filtering node identifiers prefixed with “CHEBI:” to denote biochemical entities. Records containing multiple mechanistic pathways were excluded to maintain dataset simplicity and clarity. Each qualifying pathway was formulated into the structured question: “Which biochemical entity is affected by Drug ’X’ via its mechanism of action in treating Disease ’Y’?” The metabolite node identified via CHEBI identifiers served as the ground truth answer, yielding a final dataset of 201 unique QA pairs.

Drug-Centric Benchmark: A third benchmark dataset was developed to evaluate the ability of computational models to infer therapeutic agents when provided with a disease and a mediating biological process. Pathways were selected specifically if they included exactly one BiologicalProcess node, and drugs lacking resolvable identifiers from DrugBank or MESH databases were excluded to ensure accurate and standardized identification. Each qualifying path was structured into the question: “Which drug can be used in the treatment of Disease ’Y’ by targeting Biological Process ’P’?” The corresponding drug node served as the ground truth. After thorough harmonization and stringent quality control measures, this benchmark comprised 842 unique QA pairs.

The resulting benchmarks thus offer a robust, multiscale evaluation platform specifically designed to probe the mechanistic inference capabilities of knowledge-graph-augmented language models comprehensively and rigorously.

### 2.3 Use of Large Language Models

All natural-language processing steps were carried out with two OpenAI models, GPT-4o-mini (snapshot 2024-07-18) and GPT-4o (snapshot 2024-08-06) ^32,37^. Both models were invoked through the OpenAI API. The temperature parameter was fixed at 0.0 for every request, thereby forcing deterministic decoding and facilitating reproducible evaluation. Each model accepts up to 128,000 input tokens and can return a maximum of 16,384 completion tokens. Although GPT-4o-mini is substantially smaller in parameter count than GPT-4o, both models share the same context window size, permitting a controlled comparison of model capacity while holding prompt length constant ^32,37^. At the time the experiments were executed, GPT-4o-mini was priced at 0.15 USD per million input tokens and 0.60 USD per million output tokens. The corresponding prices for GPT-4o were 2.50 USD and 10.00 USD, respectively. Model versions were pinned by explicit snapshot identifiers to eliminate the possibility of version drift during the study period. Snapshot documentation is archived at https://platform.openai.com/docs/models/gpt-4o-mini and https://platform.openai.com/docs/models/gpt-4o. ChatGPT was used to assist with grammar correction and to improve conciseness in the manuscript.

#### Prompt engineering

Each request began with a concise system prompt defining the model’s role ^38–40^. Two distinct system prompts were prepared per dataset: one for the standalone LLM baseline, and one tailored for the retrieval-augmented BTE–RAG workflow. Queries were provided directly to the model without additional contextual examples, employing a zero-shot prompting approach. To facilitate efficient and accurate downstream processing, the model was instructed to produce responses strictly in a predefined JSON format, omitting supplementary explanatory text.

## 3 Results

We developed BTE–RAG, a retrieval-augmented generation framework designed to enhance large language models (LLMs) by integrating mechanistic evidence from BioThings Explorer (BTE), a federated biomedical knowledge graph. BTE–RAG embeds structured, graph-derived context into prompts to improve mechanistic accuracy, ensure explicit provenance, and facilitate higher-order reasoning. We benchmarked the performance of BTE–RAG versus an LLM-only baseline across three distinct mechanistic reasoning tasks: gene identification, drug–metabolite interactions, and drug–biological-process relationships.

### 3.1 Mechanistic Gene Prediction

We first assessed the effect of knowledge graph augmentation on gene-level mechanistic inference using 798 curated drug–disease pairs from DrugMechDB. Queries were structured as: “Which gene plays the most significant mechanistic role in how Drug ’X’ treats or impacts Disease ’Y’?” Two models, GPT-4o and the smaller GPT-4o-mini were evaluated in two experimental conditions: (i) LLM-only, providing no additional context, and (ii) BTE–RAG, incorporating evidence retrieved via BTE.

Under the LLM-only condition, GPT-4o-mini correctly answered 407 queries (Supplementary Figure S2B), achieving an accuracy of 51%. Augmenting prompts with BTE-derived evidence markedly increased accuracy to 75.8% (Figure 2A), a substantial absolute improvement of 24.8 percentage points (Supplementary Figure S2C). The larger GPT-4o model demonstrated an accuracy of 69.8% in the baseline condition, which increased to 78.6% (627 correct answers, Supplementary Figure S3B) when supplemented with BTE context (Figure 2B), reflecting an absolute gain of 8.8 percentage points (Supplementary Figure S3C).

**Figure 2:**
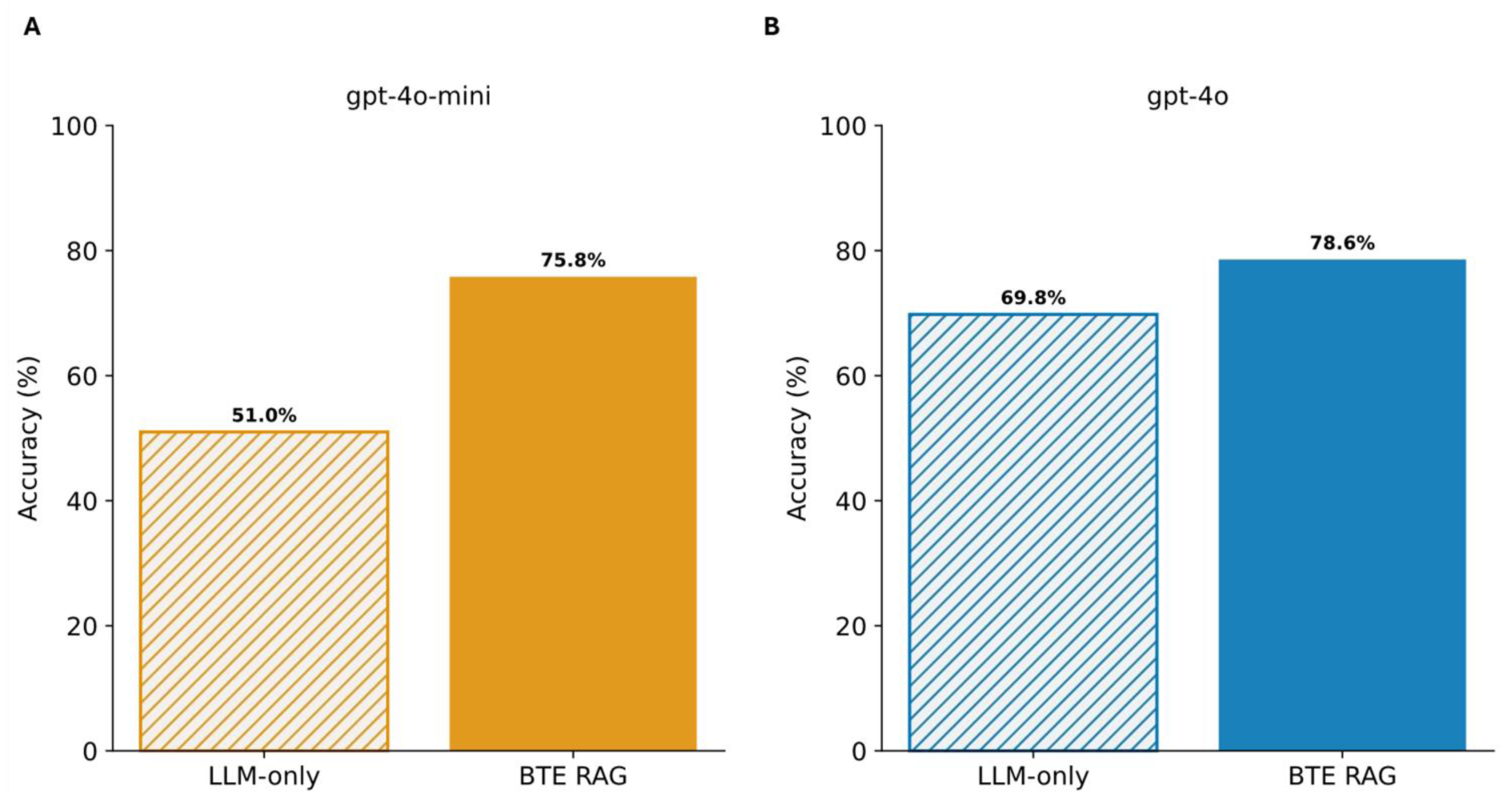
Retrieval–augmented generation with BTE-RAG markedly improves factual accuracy of gene-centric benchmark using GPT–4o models. **(A)** For the compact gpt–4o–mini model, introducing the BTE-RAG retrieval layer raised overall accuracy from 51% (hatched bar, LLM–only baseline) to 75.8 % (solid bar). **(B)** The same intervention applied to the larger gpt–4o model increased accuracy from 69.8% to 78.6 %. Accuracy was calculated as the proportion of correct answers across the composite biomedical question–answering benchmark described in Methods.

Because knowledge–graph queries can return superfluous triples, we evaluated a simple similarity–based pruning strategy. Specifically, both the user queries and the context statements were embedded using the sentence embedding model ’S-PubMedBert-MS-MARCO’ ^33^. Context statements were then ranked based on cosine similarity scores relative to the embedded query, and those statements falling within the lowest 10% similarity scores were removed to retain only the most relevant context lines. This lightweight filtering strategy preserved, and in some cases slightly enhanced performance across all evaluated accuracy metrics (Supplementary Figure S2A, S3A), suggesting that excluding the least relevant context statements can beneficially impact the accuracy of gene-level reasoning tasks. Cross-tabulation provided quantitative detail on the effect of retrieval augmentation, illustrating that BTE context flipped 245 previously incorrect answers to correct for GPT-4o-mini and 119 for GPT-4o (Supplementary Figure S2D, S3D).

Together, these findings illustrate that structured mechanistic context provided through BTE significantly enhances gene-level reasoning performance, particularly amplifying the capabilities of smaller-scale models such as GPT-4o-mini. The accuracy improvements observed even in GPT-4o highlight that state-of-the-art models retain latent knowledge gaps effectively bridged by integrating curated biomedical graphs and selectively pruning irrelevant content.

### 3.2 Prediction of Drug–Metabolite Relationships

To gauge whether retrieval augments the mechanistic fidelity of metabolite–level reasoning, we posed 201 queries of the form “Which biochemical entity is affected by Drug X via its mechanism of action in treating Disease Y?” using the DrugMechDB– derived Drug → Metabolite → Disease paths. Because metabolite names are much less standardized than gene names, we scored the answer quality by computing a semantic concordance between each model answer and the gold standard metabolite. Semantic concordance was based on cosine similarity of text embeddings using the BioBERT-STSB text embedding model, a metric that rewards graded lexical and semantic overlap rather than exact string identity ^34^.

Rank–ordered similarity curves in Figure 3A immediately reveal the effect of augmentation: for both gpt–4o–mini (orange) and gpt–4o (blue), the BTE–RAG trace (solid line) departs from the prompt–only baseline (dashed line) after ∼130 ranked questions (cosine ≈ 0.70) and widens steadily, nearly doubling the number of answers that reach the high–fidelity zone (cosine ≥ 0.90).

**Figure 3:**
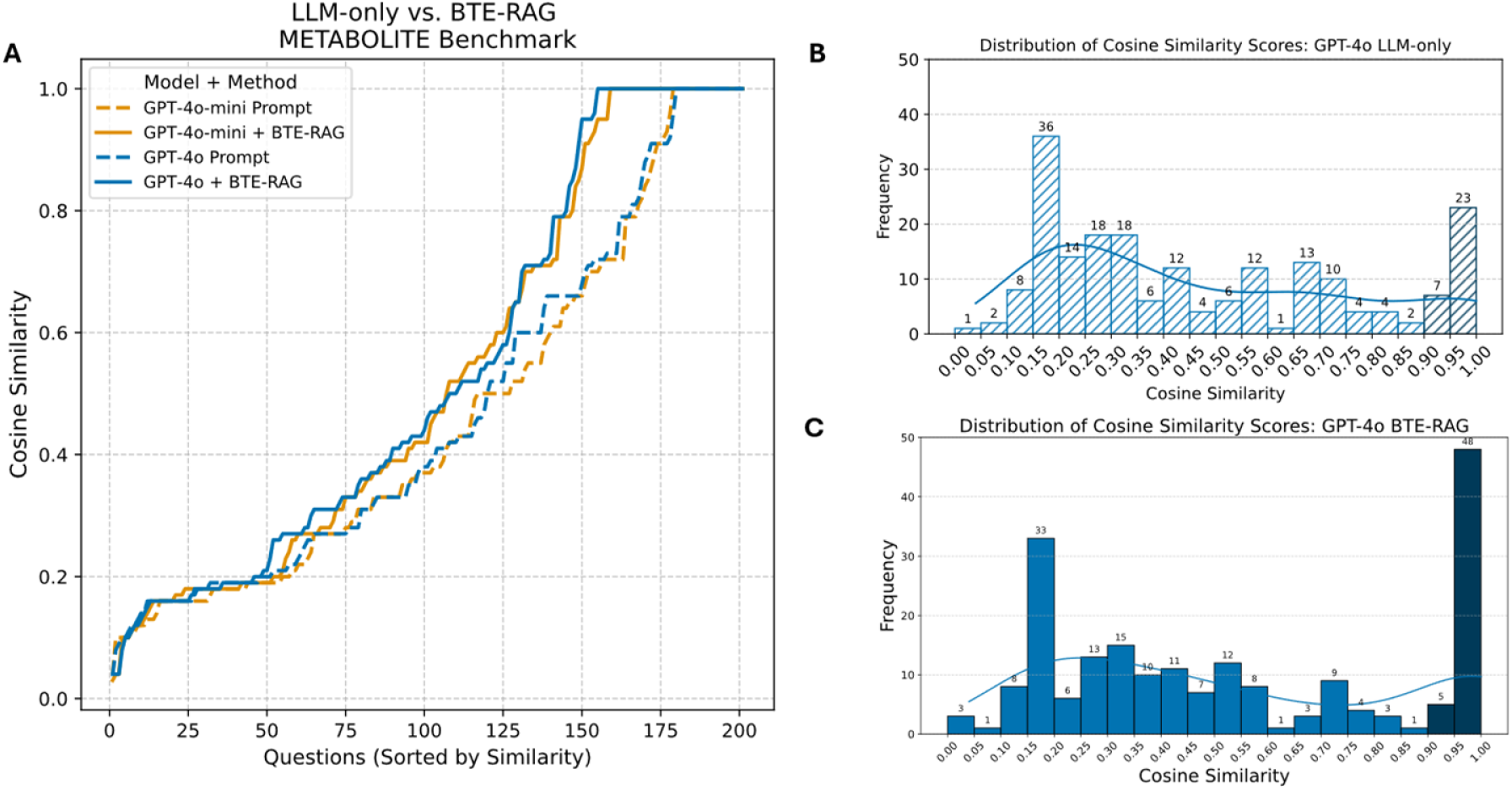
Retrieval–augmented context increases semantic concordance with ground–truth metabolites. **(A)** Cosine–similarity scores between each generated answer and the corresponding reference metabolite (sentence–transformer embeddings; see Methods) are plotted for all 201 questions in the Metabolite Benchmark, ordered from lowest to highest similarity. Dashed traces represent the LLM–only baseline, whereas solid traces include BioThings Explorer (BTE) retrieval–augmented context. Orange curves denote gpt–4o–mini; blue curves denote gpt–4o. For both model sizes, BTE–RAG systematically shifts the similarity distribution upward, indicating improved semantic alignment with the curated biochemical ground truth. **(B)** Score distribution GPT-4o, LLM-only. Histogram of cosine-similarity scores for GPT-4o answers generated without external context. Bar heights and numeric labels denote the number of questions (n = 201) falling in each bin; the overlaid KDE line summarizes the distribution. **(C)** Score distribution GPT-4o + BTE-RAG. Same format as panel B but for GPT-4o answers generated with BTE-RAG’s context. The right-shifted, more peaked distribution highlights the improvement in semantic alignment achieved by retrieval-augmented generation.

Histograms for the prompt–only condition (Figure 3B, gpt4o; Supplementary Figure S4, gpt-4o-mini) reveal a pronounced left–skew: both gpt–4o–mini and gpt–4o peak in the 0.15–0.30 similarity bins, with medians below 0.30. Only 15 % of answers fall in the high–similarity regime (≥ 0.90), indicating that the LLMs frequently retrieve metabolites that are semantically distant from the curated ground truth.

Appending BTE evidence shifts the distributions rightward across similarity bins (Figure 3C (gpt-4o), Supplementary Figure S5, S6). For GPT-4o-mini, applying a stringent context similarity threshold (>80th percentile) increased the number of high-fidelity answers (cosine similarity 0.90–1.00) from 28 to 51 (+82%). Similarly, GPT-4o exhibited an increase from 30 to 53 (+77%) under the same conditions. Simultaneously, counts in the mid–similarity interval (0.40–0.70) contract (Supplementary Figure S5, S6), confirming that retrieval largely converts borderline predictions into highly concordant hits rather than merely redistributing low–score failures.

Because voluminous context can inflate token budgets, we assessed performance when progressively discarding lower–ranked context lines (10th to 90th percentile cut–offs). Rank–ordered similarity traces (Supplementary Figure S7) show that the BTE–RAG curves remain above or coincide with the prompt–only baseline throughout the distribution even when 90 % of context is withheld. Histograms (Supplementary Figure S5, S6) reinforce this observation: the ≥ 0.90 similarity bin consistently retains ≥ 40 hits for both models across all pruning levels, demonstrating that a concise subset of top–ranked evidence lines is sufficient to drive the bulk of the performance gains.

### 3.3 Drug–Biological Process Reasoning

We next asked 842 DrugMechDB questions of the form “Which drug can be used in the treatment of Disease Y by targeting Biological Process P?”. Answer fidelity was again scored with BioBERT–STSB cosine similarity ^34^. In rank–ordered plots (Figure 4A), the prompt–only (dashed) and BTE–RAG (solid) curves for both gpt–4o–mini (orange) and gpt–4o (blue) are nearly super–imposable through the first ≈ 600 ranked queries (cosine < 0.70). Beyond this inflection point, the BTE–augmented traces bend upward more steeply, yielding a clear margin in the high–fidelity zone (cosine ≥ 0.80). Thus, retrieval does not alter overall parity but selectively boosts the most mechanistically demanding subset of questions.

**Figure 4:**
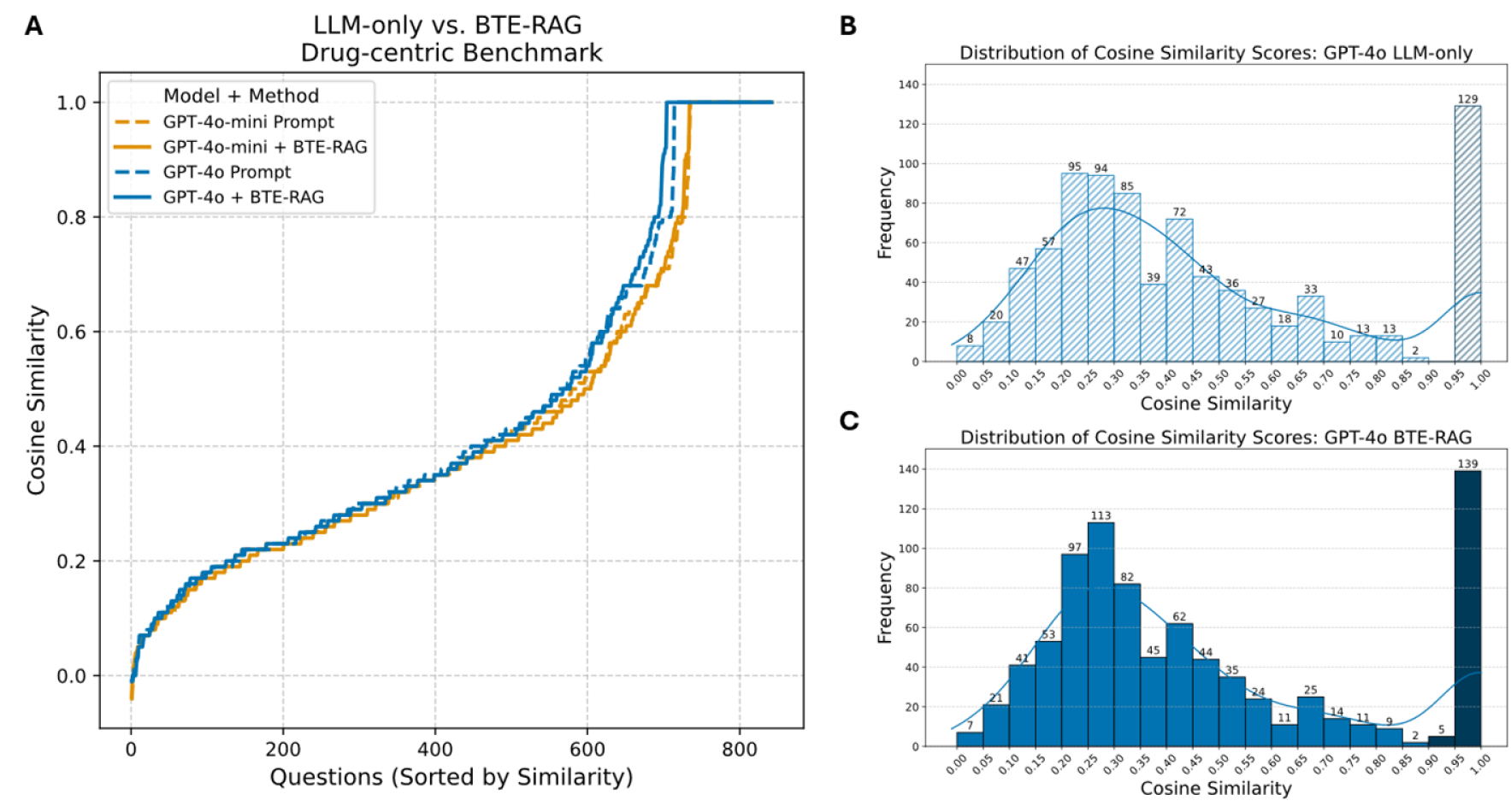
Retrieval-augmented generation maintains overall parity yet excels in the high-fidelity regime of drug-centric mechanistic answers. **(A)** Cosine-similarity scores (sentence-transformer embeddings; see *Methods*) between each generated answer and the reference drug→biological-process pathway are plotted for all 842 questions in the Drug Benchmark, ordered from lowest to highest similarity. Dashed traces (LLM-only) and solid traces (BTE-RAG) follow nearly overlapping trajectories across most of the distribution, indicating broadly comparable performance between the two inference modes. However, above a cosine similarity threshold of ≈ 0.7, both *gpt-4o-mini* (orange) and *gpt-4o* (blue) curves generated with BTE context surge ahead of their prompt-only counterparts, revealing a marked advantage in producing highly concordant mechanistic explanations. **(B)** Score distribution GPT-4o, LLM-only. Histogram of cosine-similarity scores for GPT-4o answers generated without external context. The hatched bar at 0.90–1.00 marks the high-fidelity zone, capturing 129 near-perfect matches produced by the baseline model. **(C)** Score distribution GPT-4o + BTE-RAG. Same format as panel B but for GPT-4o answers produced with BTE-RAG’s context. The distribution is right-shifted, and the solid bar in the 0.90–1.00 high-fidelity zone now contains 144 answers, highlighting the enrichment of top-tier mechanistic concordance achieved through retrieval-augmented generation.

Prompt–only histograms (Figure 4B; Supplementary Figure S8, gpt-4o-mini) peak in the 0.20–0.35 range, with ∼15 % of answers falling in the ≥ 0.90 bin. Appending the full BTE context nudges the entire distribution rightward (Figure 4C; Supplementary Figure S9-top–left panels). The ≥ 0.90 bin increases by ≈ 5–10 % for both model sizes. These shifts, though smaller than those seen for gene– and metabolite tasks, account for the late–stage separation observed in Figure 4A.

Unlike the previous tasks, performance here depends on retaining a broad evidentiary window. When the lowest–ranked 10–20 % of context lines are removed, the uplift in the ≥ 0.90 bin attenuates, and the rank–ordered curves progressively converge toward the baseline (Supplementary Figure S9, S10 & S11). Deeper cuts (> 40 %) essentially erase the retrieval advantage. This suggests that pathway–level questions draw on a more diffuse set of graph triples than gene or metabolite queries, and aggressive trimming can discard critical relational clues. For drug → biological–process reasoning, BTE–RAG delivers targeted gains in the top decile of similarity scores, provided the complete knowledge–graph context is supplied.

These findings reinforce that optimal evidence granularity is task–dependent: concise, high–relevance snippets suffice for gene– and metabolite–level inference, whereas pathway–level queries benefit from a richer contextual fabric. By grounding LLM outputs within curated, biologically meaningful pathways, BTE–RAG consistently accelerates accurate inference, reduces residual errors, and demonstrates considerable promise for advancing automated biomedical hypothesis generation and therapeutic repurposing workflows.

## 4 Discussion

The rapid advancement of large language models (LLMs) has profoundly reshaped biomedical natural language processing ^41^. Despite these advances, current LLMs predominantly operate as opaque systems with implicit knowledge representation, rendering their factual accuracy challenging to verify and limiting their applicability in high-stakes biomedical environments. Recent efforts, such as the knowledge-graph augmented retrieval approach ^21^, have successfully enhanced biomedical reasoning by integrating disease-specific embeddings from specialized knowledge graphs such as SPOKE ^42^. We developed BTE–RAG, a novel retrieval-augmented generation pipeline that strategically incorporates explicit mechanistic evidence from BTE ^27^. By leveraging the extensive and federated biomedical knowledge graph of BTE, our method substantially broadens the applicability of knowledge-graph augmented strategies to address diverse query types, including those involving genes, proteins, metabolites, biological processes, diseases and chemical substances. This capability allows BTE– RAG to support complex, multi-domain biomedical inquiries, significantly extending beyond disease-centric queries alone. Our comparative analysis, utilizing a direct “LLM-only” approach versus the BTE-augmented strategy (Figure 1A) across three rigorously constructed DrugMechDB benchmarks (Figure 1B), demonstrates that incorporating explicit, structured context significantly elevates answer accuracy, enhances transparency, and allows smaller, more computationally efficient models to perform competitively with leading-edge systems. The granularity, explicit mechanistic grounding, and high-quality source attribution inherent in these benchmarks uniquely position them for probing the causal inference capabilities of language models. Comparable mechanistically focused datasets remain scarce in the biomedical domain, as existing resources like PubMedQA or Natural Questions predominantly target document-level retrieval or summarization rather than deep mechanistic inference ^43,44^.

Traditional LLMs accumulate domain-specific knowledge implicitly during pre-training by statistically modeling large collections of biomedical texts. Although this method yields linguistically coherent responses, it inherently exposes models to the risk of hallucinations, particularly in scenarios involving sparse biomedical facts or multi-step mechanistic reasoning. By contrast, retrieval-augmented generation explicitly anchors model predictions in verifiable external sources, constraining generation to well-substantiated evidence. BTE–RAG advances this paradigm by dynamically federating 61 authoritative biomedical APIs into a single cohesive meta-graph, thereby enabling real-time inclusion of newly curated knowledge in generated responses and ensuring reproducible benchmarking through cached retrievals.

Four critical design principles underpin the efficacy of the BTE–RAG framework. First, the framework leverages an API-centric federation layer that integrates trusted biomedical data sources, including MyGene.info, Gene Ontology, CTDbase, Pubmed central, CHEBI, disease-ontology, DrugBank and more, through unified interface of BTE ^29,36,45–47^. Second, it employs semantic query templates aligned with the Translator Reasoner API (TRAPI) standard, selectively retrieving only the most relevant relationships for each question, thereby avoiding extraneous contextual noise. Third, retrieved knowledge graph triples are translated into succinct, directionally explicit declarative statements, seamlessly integrating structured knowledge with natural-language prompts. Fourth, BTE–RAG incorporates flexible context-selection strategies; full-context utilization and cosine similarity-based pruning for scenarios requiring concise, highly relevant context subsets.

Across diverse mechanistic tasks, including gene-centric, metabolite-centric, and drug-centric benchmarks derived from DrugMechDB ^31^, BTE-augmented prompting consistently outperformed the LLM-only approach. Notably, the smaller GPT-4o-mini model achieved over sixty-percentage improvement in accuracy on the gene-centric task and eighty-two percent improvement on the metabolite task, when provided with structured BTE evidence. Even GPT-4o, the larger flagship model, demonstrated substantial accuracy gains, underscoring that high-quality, explicit mechanistic context can effectively mitigate the need for extremely large model sizes, suggesting a cost-efficient pathway toward domain-specific accuracy.

While BTE offers comprehensive coverage across numerous biomedical domains, certain areas such as single-cell data, epigenomic profiles, and microbiome interactions remain sparsely represented. Furthermore, variations in curation quality across federated APIs could inadvertently propagate erroneous edges into model-generated contexts. Although our evaluation leveraged the meticulously curated, high-confidence knowledge graph of DrugMechDB, real-world applications may require strategies for managing lower-confidence or conflicting evidence. Our study employed deterministic prompting to maintain comparability; exploring guided, chain-of-thought prompting strategies could further enhance complex reasoning capabilities but may simultaneously reintroduce hallucinatory risks.

Future developments of BTE–RAG may involve integration into autonomous agent systems capable of iterative querying, generation, self-critiquing, and re-querying, thus facilitating automated self-verification workflows. Expanding the underlying knowledge graph to incorporate resources such as LINCS transcriptomic signatures, tissue-specific interaction networks, and multi-omics datasets would further enrich the mechanistic coverage and broaden applicability ^48^. Expanding benchmarking efforts beyond DrugMechDB to encompass open-world biomedical queries could rigorously evaluate and strengthen the capacity BTE–RAG for reliable, contextually grounded inference. Furthermore, adopting frameworks like the Model Context Protocol could harmonize comparisons across diverse generative models, facilitate rigorous auditing, and support real-time decision-making in clinical and regulatory contexts.

In conclusion, BTE–RAG demonstrates the substantial value derived from strategically integrating explicit mechanistic evidence into biomedical language modeling workflows. By significantly improving answer accuracy, interpretability, and computational efficiency, this approach provides a scalable, transparent, and robust foundation for future biomedical AI systems, effectively balancing accuracy, affordability, and trustworthiness.

## Supplementary File

**Supplementary Figure S1:** Detailed pipeline for BTE-RAG

**Supplementary Figure S2:** Performance of BTE–RAG versus an LLM–only baseline on the gene–centric benchmark using gpt–4o–mini.

**Supplementary Figure S3:** Performance of BTE–RAG versus an LLM–only baseline on the gene–centric benchmark using gpt–4o.

**Supplementary Figure S4:** Cosine-similarity profile for the metabolite-centric benchmark using GPT-4o-mini in LLM-only mode.

**Supplementary Figure S5:** Distribution of answer similarities for the metabolite-centric benchmark using GPT-4o-mini in BTE-RAG mode.

**Supplementary Figure S6:** Distribution of answer similarities for the metabolite-centric benchmark using GPT-4o in BTE-RAG mode.

**Supplementary Figure S7:** Rank-ordered cosine similarities between model predictions and ground-truth answers on the metabolite-centric benchmark, across context filtering thresholds.

**Supplementary Figure S8:** Cosine-similarity profile for the drug-centric benchmark using GPT-4o-mini in LLM-only mode.

**Supplementary Figure S9:** Distribution of answer similarities for the drug-centric benchmark using GPT-4o-mini in BTE-RAG mode.

**Supplementary Figure S10:** Distribution of answer similarities for the drug-centric benchmark using GPT-4o in BTE-RAG mode.

**Supplementary Figure S11:** Rank-ordered cosine similarities between model predictions and ground-truth answers on the drug-centric benchmark, across context filtering thresholds.

**Supplementary Table S1**: System prompts used for each task and model

## Availability of Data and Materials

The source code, datasets, and analysis workflows described in this manuscript are publicly available in the GitHub repository: Project Name: BTE-RAG. Repository URL: https://github.com/janjoy/BTE-RAG

## Author contributions

J.J. and A.S. conceived the project and proposed the benchmark curation. J.J. implemented the code, created the benchmarks and wrote the manuscript. A.S. supervised the study. All authors read and approved the final manuscript.

## Acknowledgements

We thank Jackson Callaghan, Mikhael Astorga, and Karthik Soman for insightful discussions, and Everaldo Rodolpho for technical support with high-performance computing resources and server infrastructure.

## Supplementary Figures

**Figure S1:**
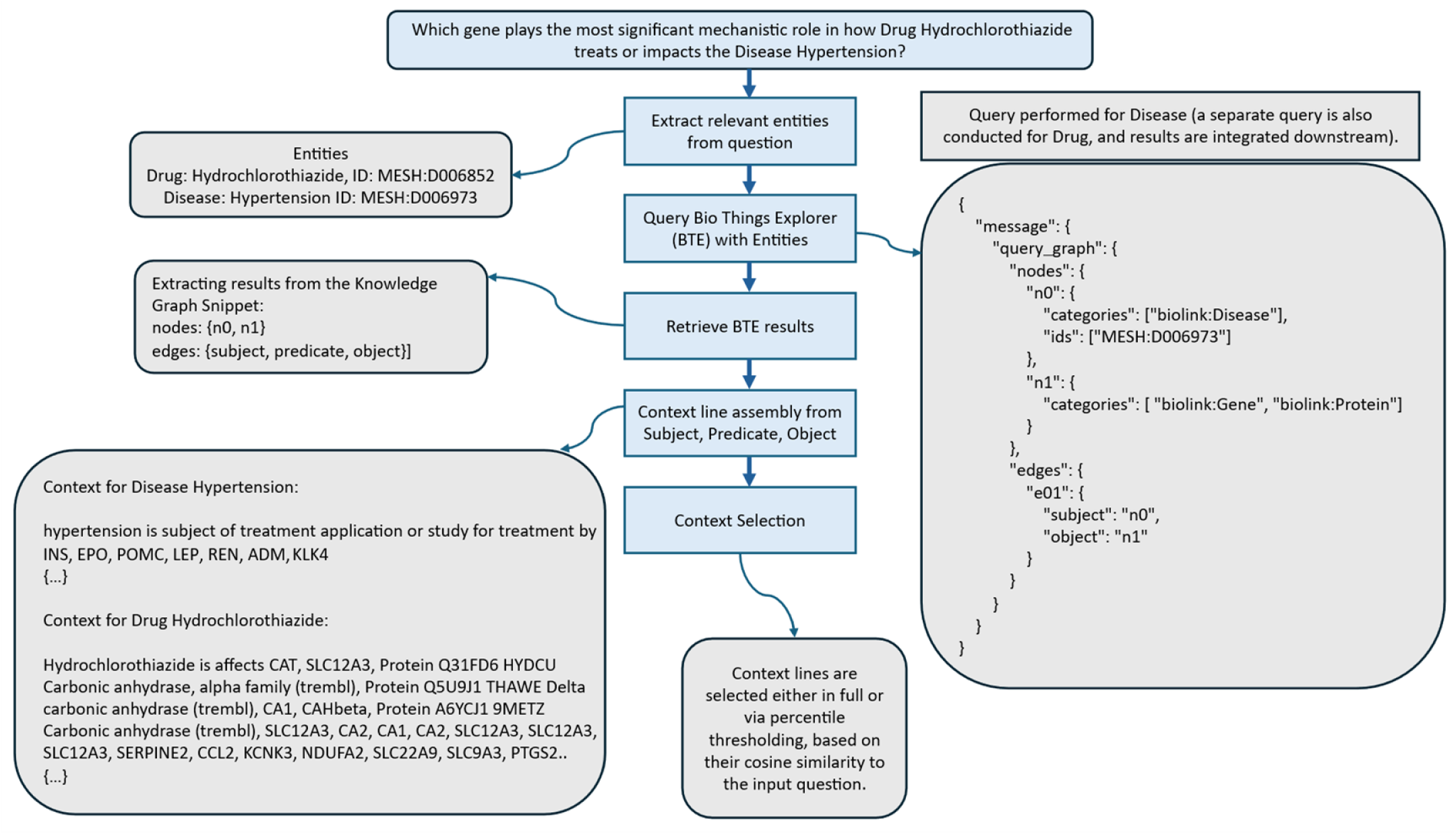
Detailed pipeline for BTE-RAG. Supplementary Figure S1 depicts the end-to-end workflow through which the BTE-RAG retrieval module converts a biomedical question into the evidence snippets ultimately supplied to the language-model reasoner. Beginning with an example query, “Which gene plays the most significant mechanistic role in how the drug *hydrochlorothiazide* treats or impacts the disease *hypertension*?”, the system first performs named-entity recognition, normalizing the detected concepts to controlled identifiers (Drug: MESH:D006852; Disease: MESH:D006973). Each entity is then submitted to BioThings Explorer (BTE) as part of a query graph that requests mechanistically relevant genes and proteins; independent queries are executed for the drug and for the disease. BTE returns knowledge-graph sub-graphs whose nodes and edges represent subject-predicate-object triples grounded in the biomedical literature. These triples are linearized into plain-text sentences, yielding two preliminary corpora (one for the disease, one for the drug) that list, for example, genes such as *INS*, *REN*, *SLC12A3* and *PTGS2* with their associated predicates. Finally, the complete set of sentences or a percentile-filtered subset is ranked by cosine similarity to the original question, and the highest-scoring lines are selected as the “retrieved context” passed forward for answer generation.

**Figure S2:**
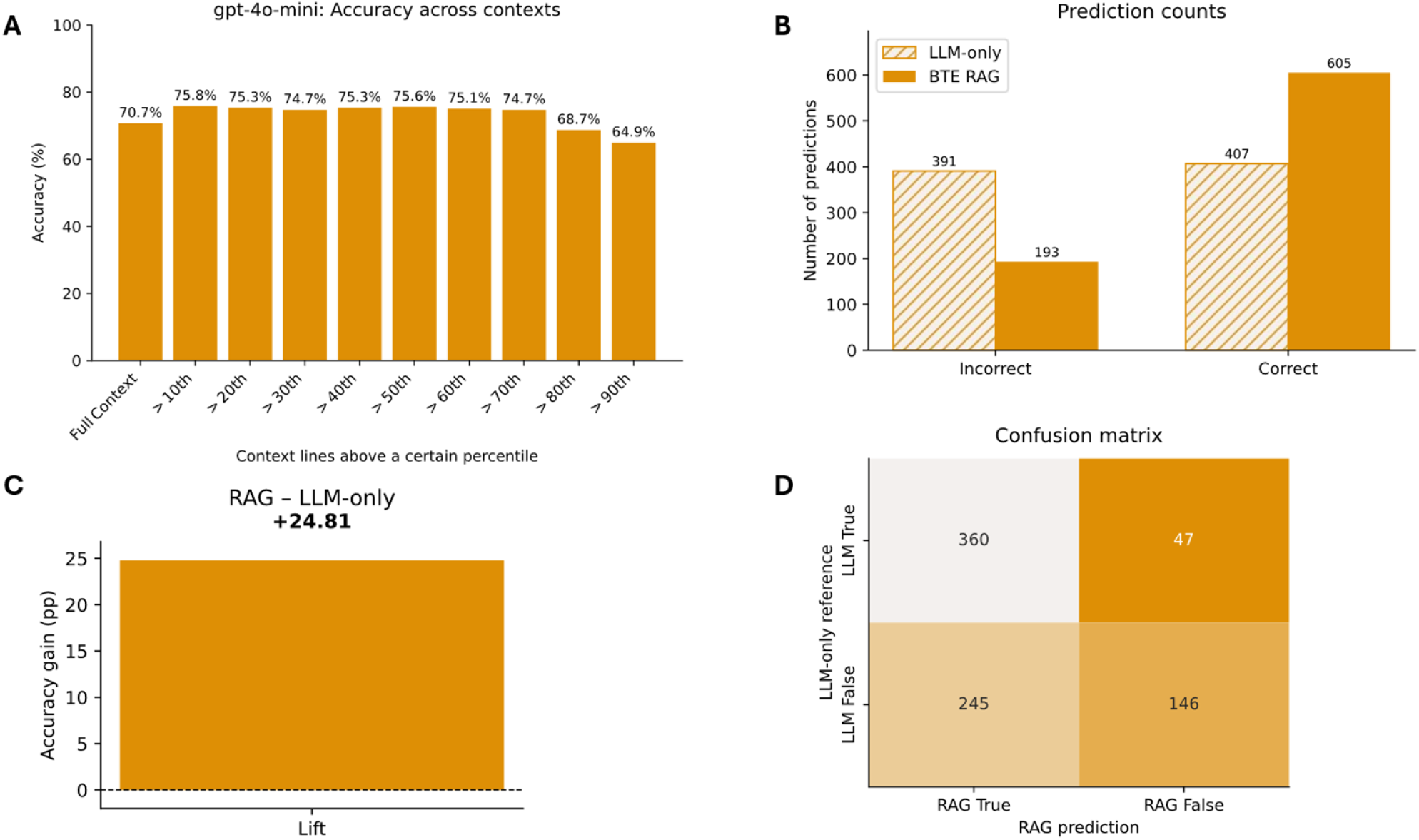
Performance of BTE–RAG versus an LLM–only baseline on the gene–centric benchmark using gpt–4o–mini. (A) Overall accuracy as a function of how much of the retrieved context is retained. Bars show accuracy when only context lines above a given cosine–similarity percentile are supplied to the model (10 th–90 th) as well as when the full context is used. (B) Breakdown of prediction counts for the 798 benchmark questions. The hatched bars represent the LLM–only condition; solid bars represent BTE–RAG. (C) BTE–RAG outperforms the LLM–only run by +24.8 percentage points, confirming that targeted knowledge–graph snippets materially improve answer quality. Confusion matrix comparing the two methods. The upper–left cell (360 cases) denotes questions both methods answer correctly; the lower–left cell (245) highlights errors that BTE–RAG fixes; the upper–right cell (47) shows instances where retrieval introduces an error; and the lower–right cell (146) comprises questions neither approach resolves.

**Figure S3:**
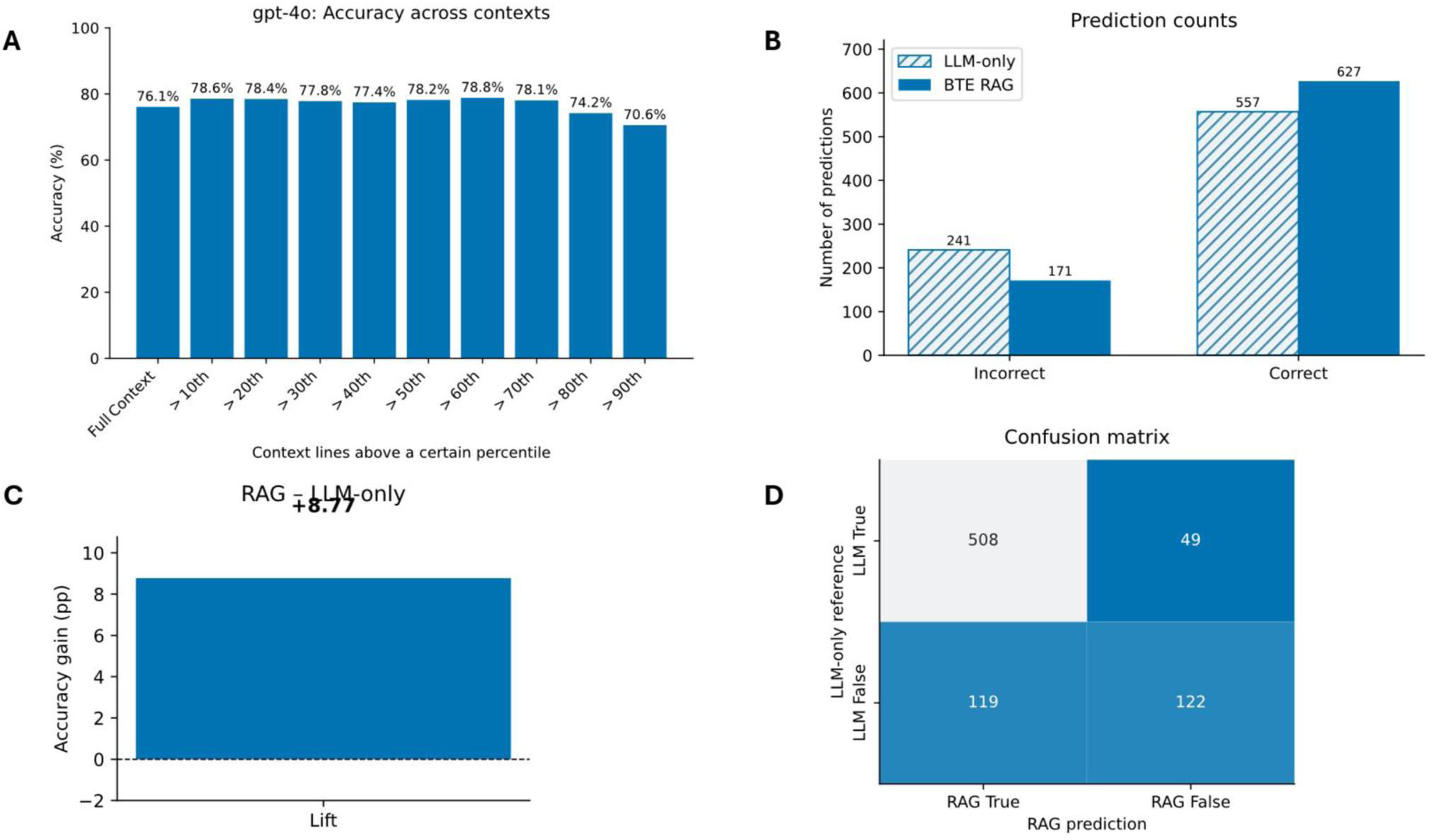
Performance of BTE–RAG versus an LLM–only baseline on the gene–centric benchmark using gpt–4o. (A) Overall accuracy as a function of how much of the retrieved context is retained. Bars show accuracy when only context lines above a given cosine–similarity percentile are supplied to the model (10 th–90 th) as well as when the full context is used. (B) Breakdown of prediction counts for the 798 benchmark questions. The hatched bars represent the LLM–only condition; solid bars represent BTE–RAG. (C) BTE–RAG outperforms the LLM–only run by +8.8 percentage points, confirming that targeted knowledge–graph snippets materially improve answer quality. Confusion matrix comparing the two methods. The upper–left cell (508 cases) denotes questions both methods answer correctly; the lower–left cell (119) highlights errors that BTE–RAG fixes; the upper–right cell (49) shows instances where retrieval introduces an error; and the lower–right cell (122) comprises questions neither approach resolves.

**Figure S4:**
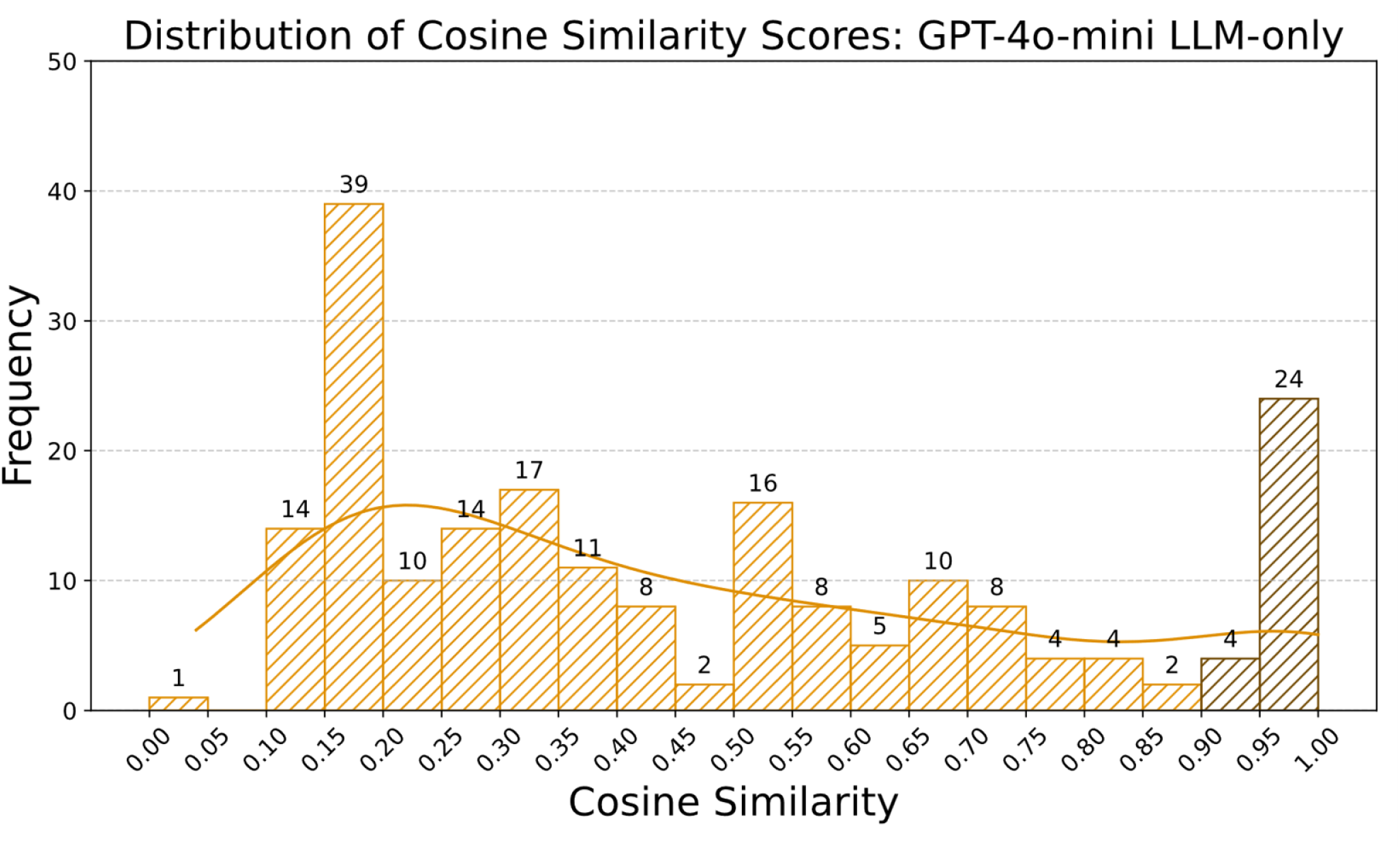
Cosine-similarity profile for the metabolite-centric benchmark using GPT-4o-mini in LLM-only mode. Histogram shows the frequency of cosine-similarity scores (bin width = 0.05) between model answers and ground truth answers across 201 metabolite-related queries when using gpt-4o-mini. A smoothed kernel-density curve traces the overall score profile.

**Figure S5:**
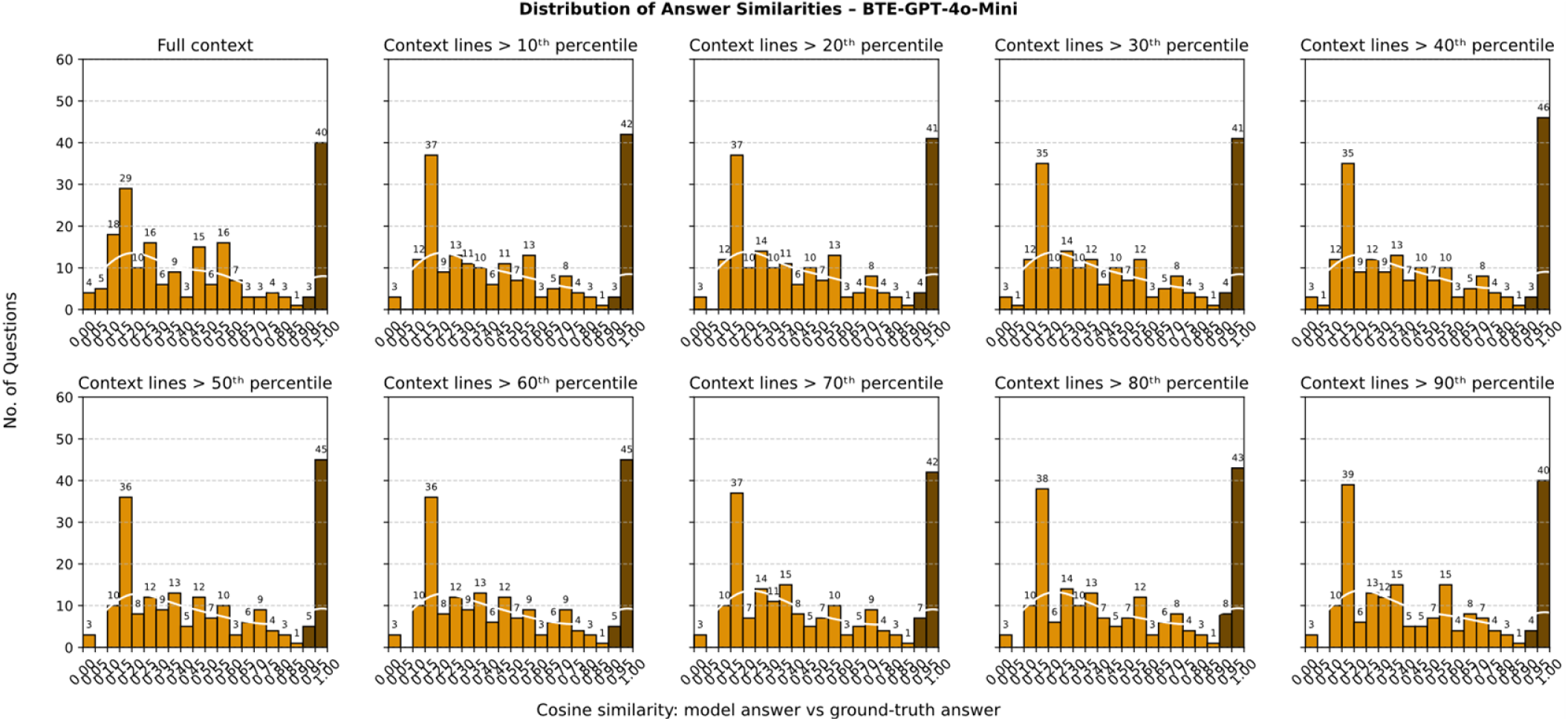
Distribution of answer similarities for the metabolite-centric benchmark using GPT-4o-mini in BTE-RAG mode. Each panel shows the cosine similarity between model predictions and ground-truth answers when either the full retrieved context is used (top left) or when context lines are filtered above increasing cosine similarity percentiles (10th to 90th).

**Figure S6:**
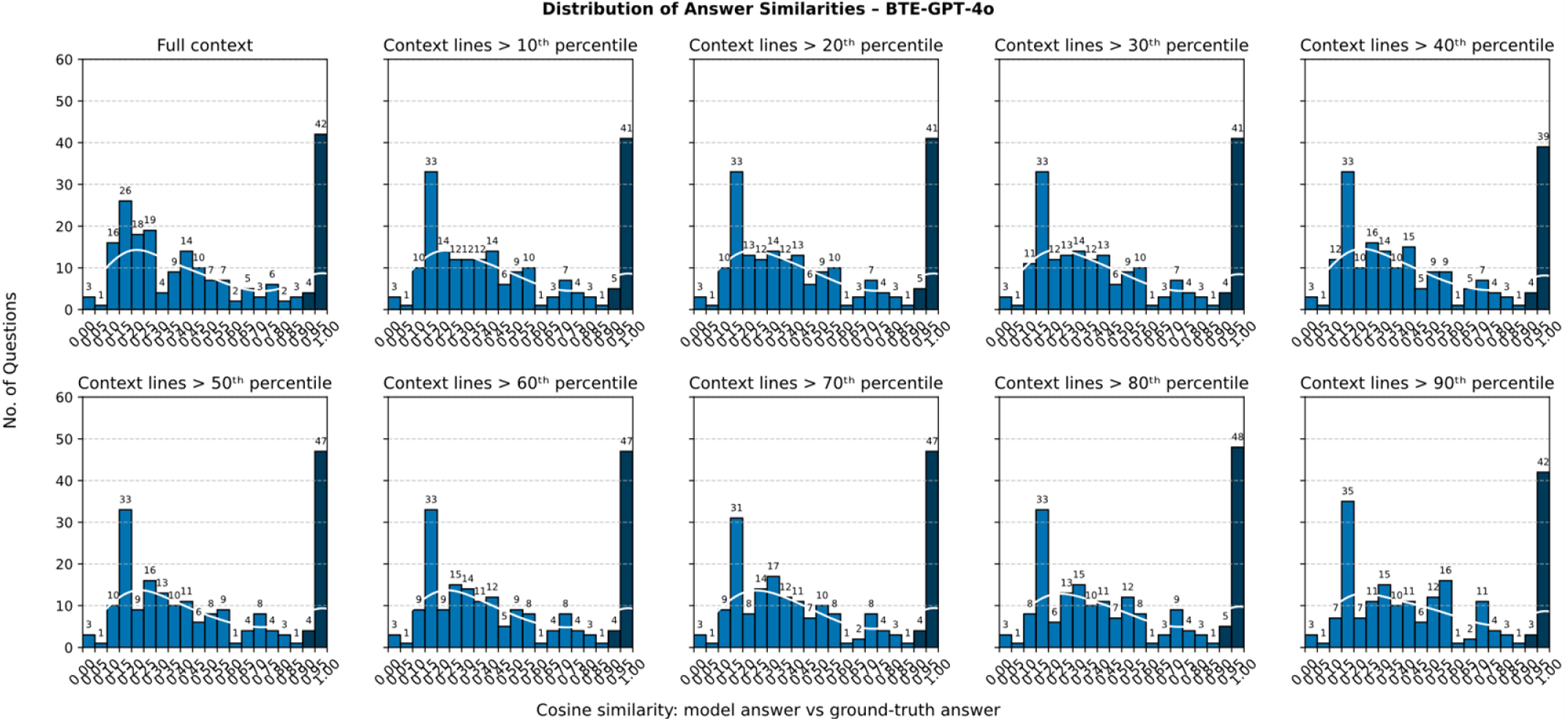
Distribution of answer similarities for the metabolite-centric benchmark using GPT-4o in BTE-RAG mode. Each panel shows the cosine similarity between model predictions and ground-truth answers when either the full retrieved context is used (top left) or when context lines are filtered above increasing cosine similarity percentiles (10th to 90th).

**Figure S7:**
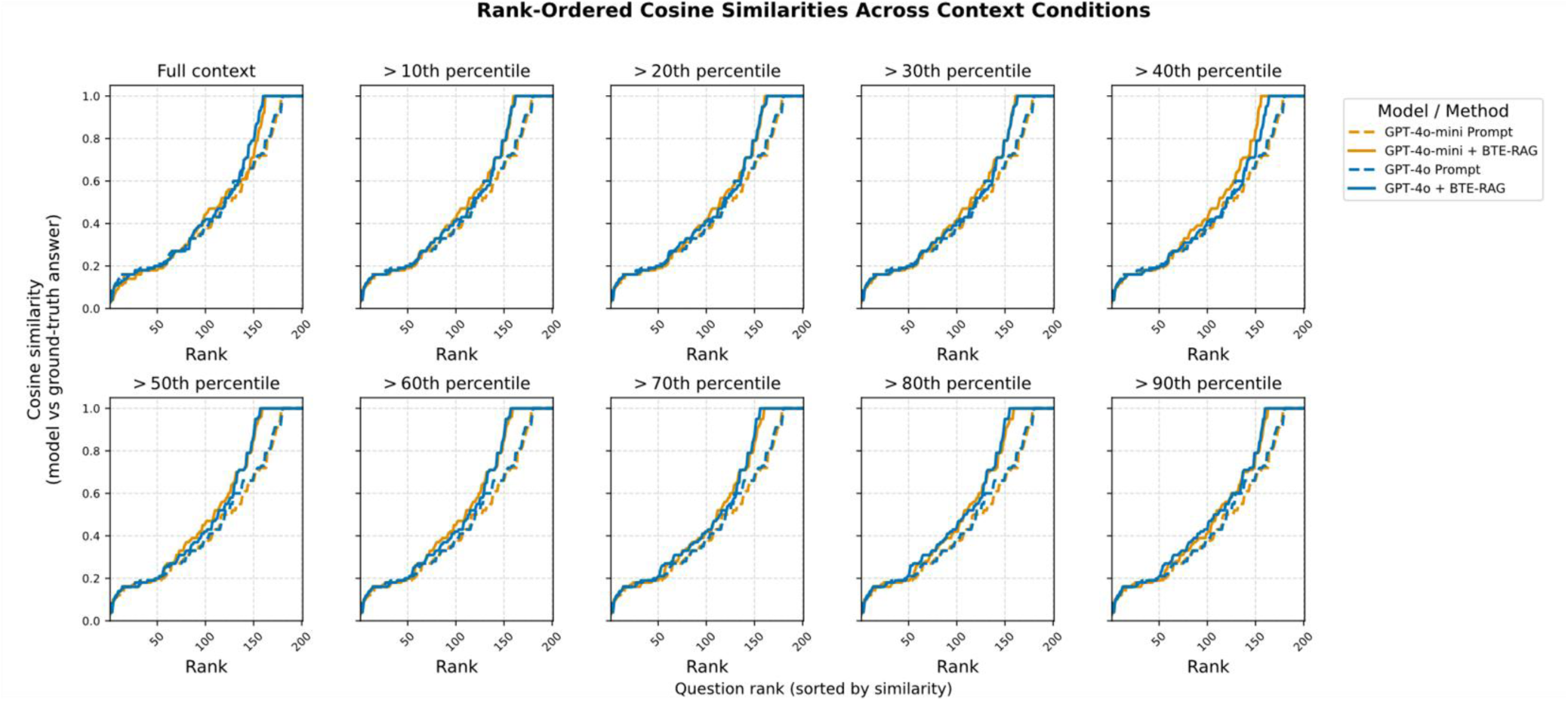
Rank-ordered cosine similarities between model predictions and ground-truth answers on the metabolite-centric benchmark, across context filtering thresholds. Each panel displays results from four model–method combinations (GPT-4o-mini-Prompt (LLM-only), GPT-4o-mini + BTE-RAG, GPT-4o Prompt, GPT-4o + BTE-RAG) under either full context or filtered context lines exceeding the indicated cosine similarity percentile (10th to 90th). Question predictions are sorted by similarity, revealing how context filtering and model selection affect semantic alignment with ground truth.

**Figure S8:**
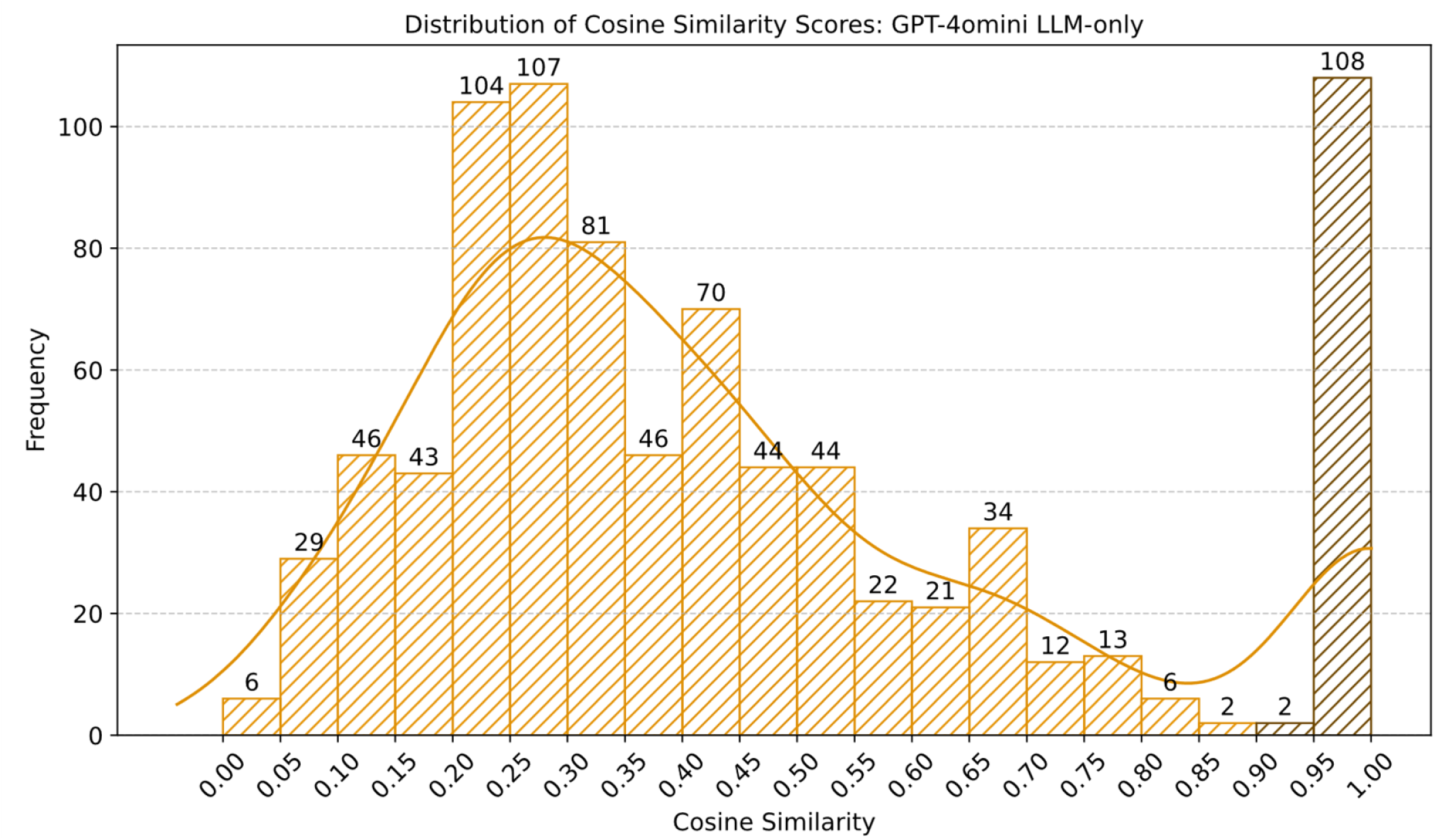
Cosine-similarity profile for the drug-centric benchmark using GPT-4o-mini in LLM-only mode. Histogram shows the frequency of cosine-similarity scores (bin width = 0.05) between model answers and ground truth answers across 842 drug-biological process queries when using gpt-4o-mini. A smoothed kernel-density curve traces the overall score profile.

**Figure S9:**
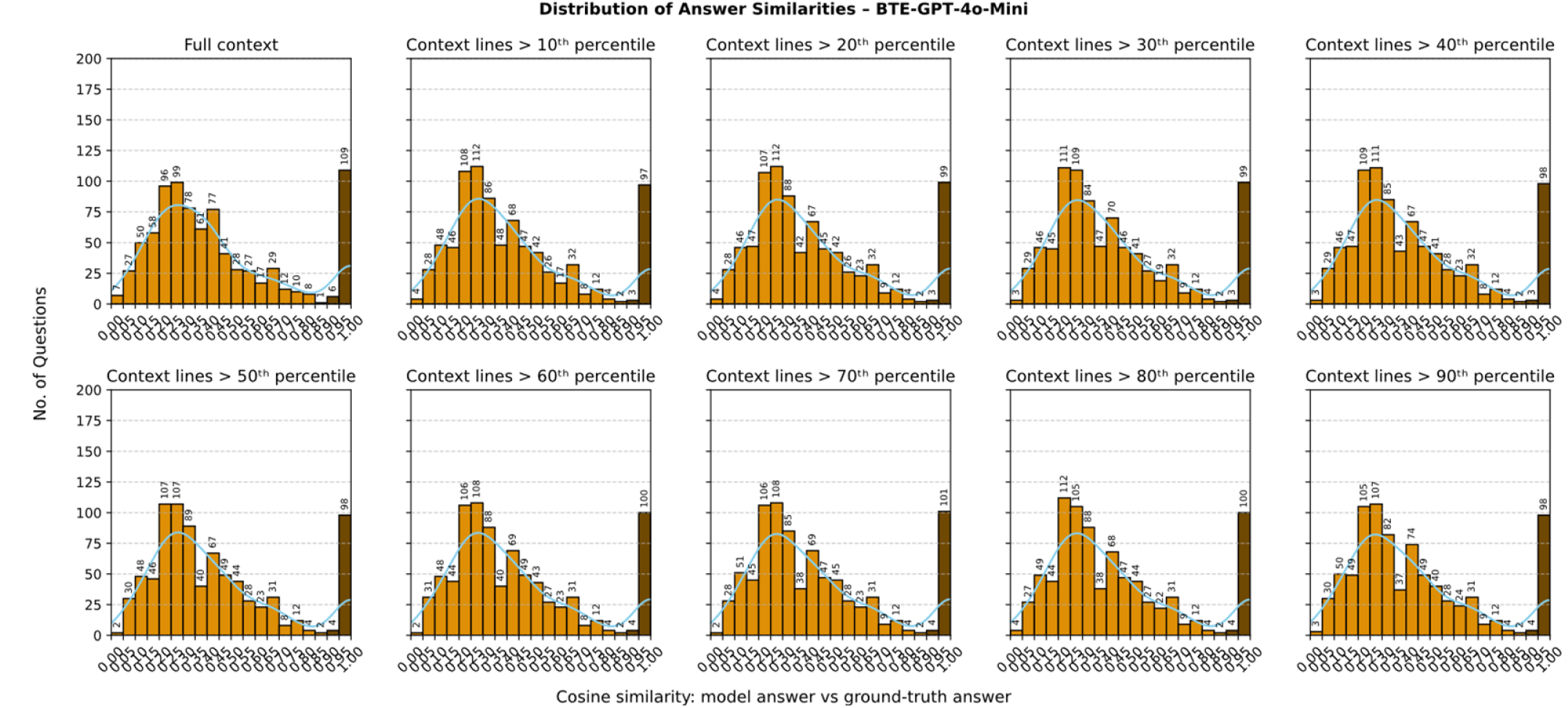
Distribution of answer similarities for the drug-centric benchmark using GPT-4o-mini in BTE-RAG mode. Each panel shows the cosine similarity between model predictions and ground-truth answers when either the full retrieved context is used (top left) or when context lines are filtered above increasing cosine similarity percentiles (10th to 90th).

**Figure S10:**
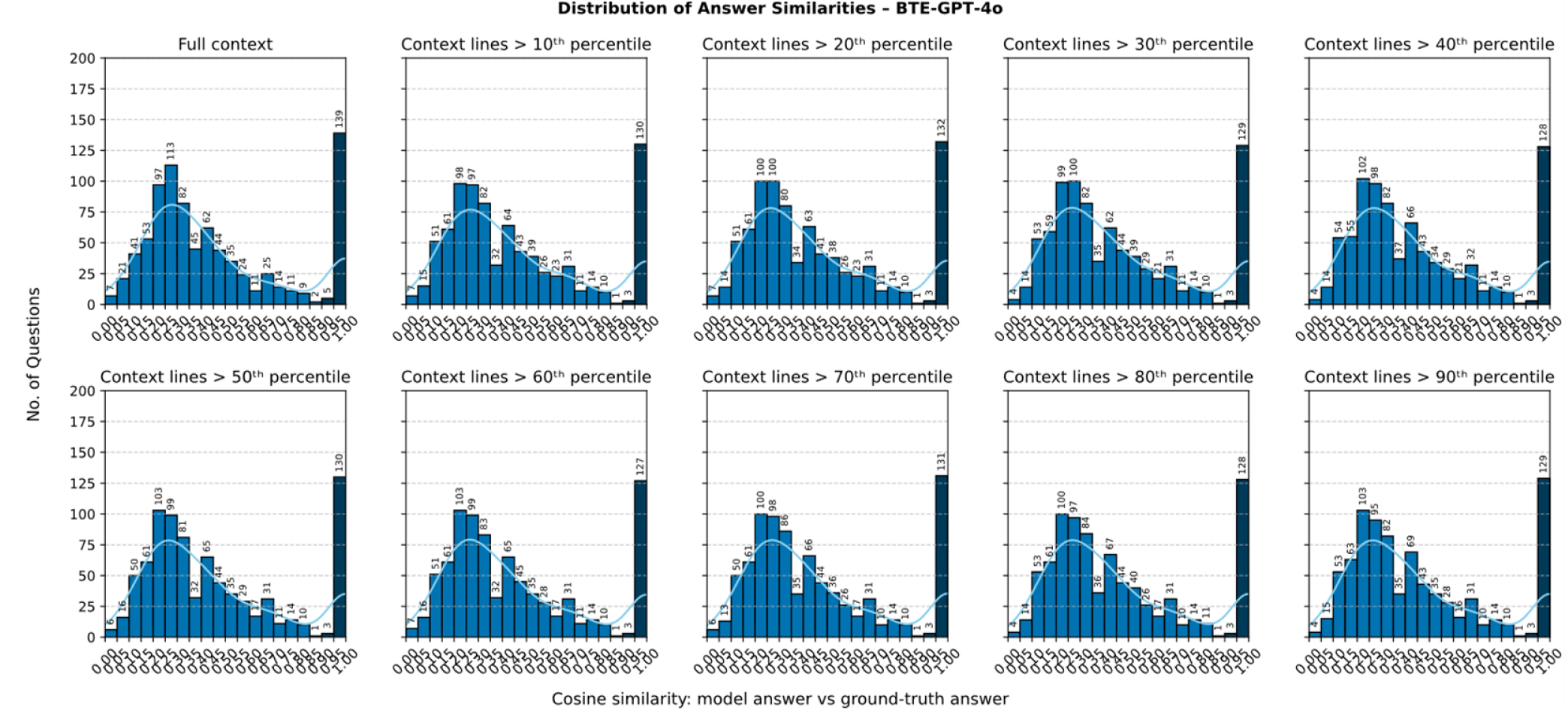
Distribution of answer similarities for the drug-centric benchmark using GPT-4o in BTE-RAG mode. Each panel shows the cosine similarity between model predictions and ground-truth answers when either the full retrieved context is used (top left) or when context lines are filtered above increasing cosine similarity percentiles (10th to 90th).

**Figure S11:**
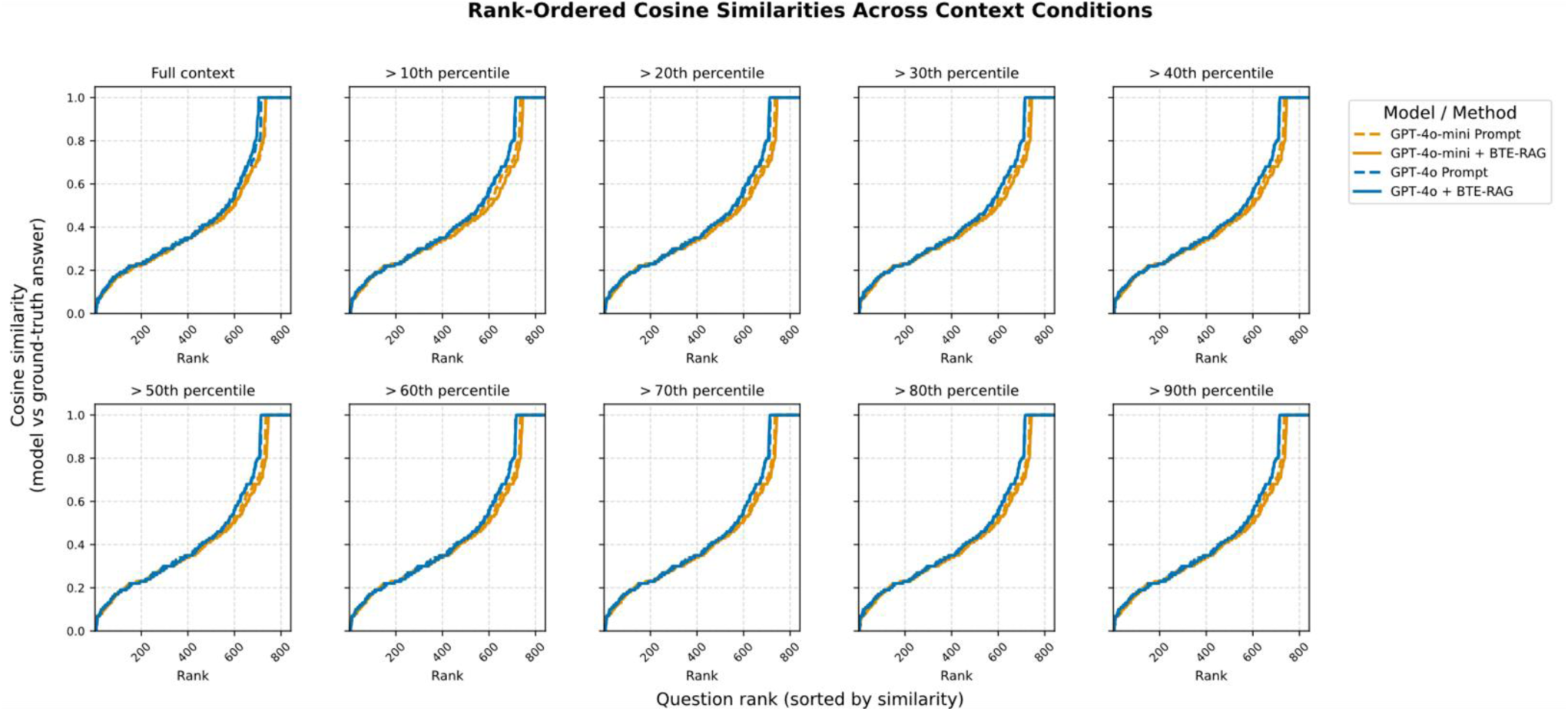
Rank-ordered cosine similarities between model predictions and ground-truth answers on the drug-centric benchmark, across context filtering thresholds. Each panel displays results from four model–method combinations (GPT-4o-mini Prompt (LLM-only), GPT-4o-mini + BTE-RAG, GPT-4o Prompt, GPT-4o + BTE-RAG) under either full context or filtered context lines exceeding the indicated cosine similarity percentile (10th to 90th). Question predictions are sorted by similarity, revealing how context filtering and model selection affect semantic alignment with ground truth.

**Table S1:**
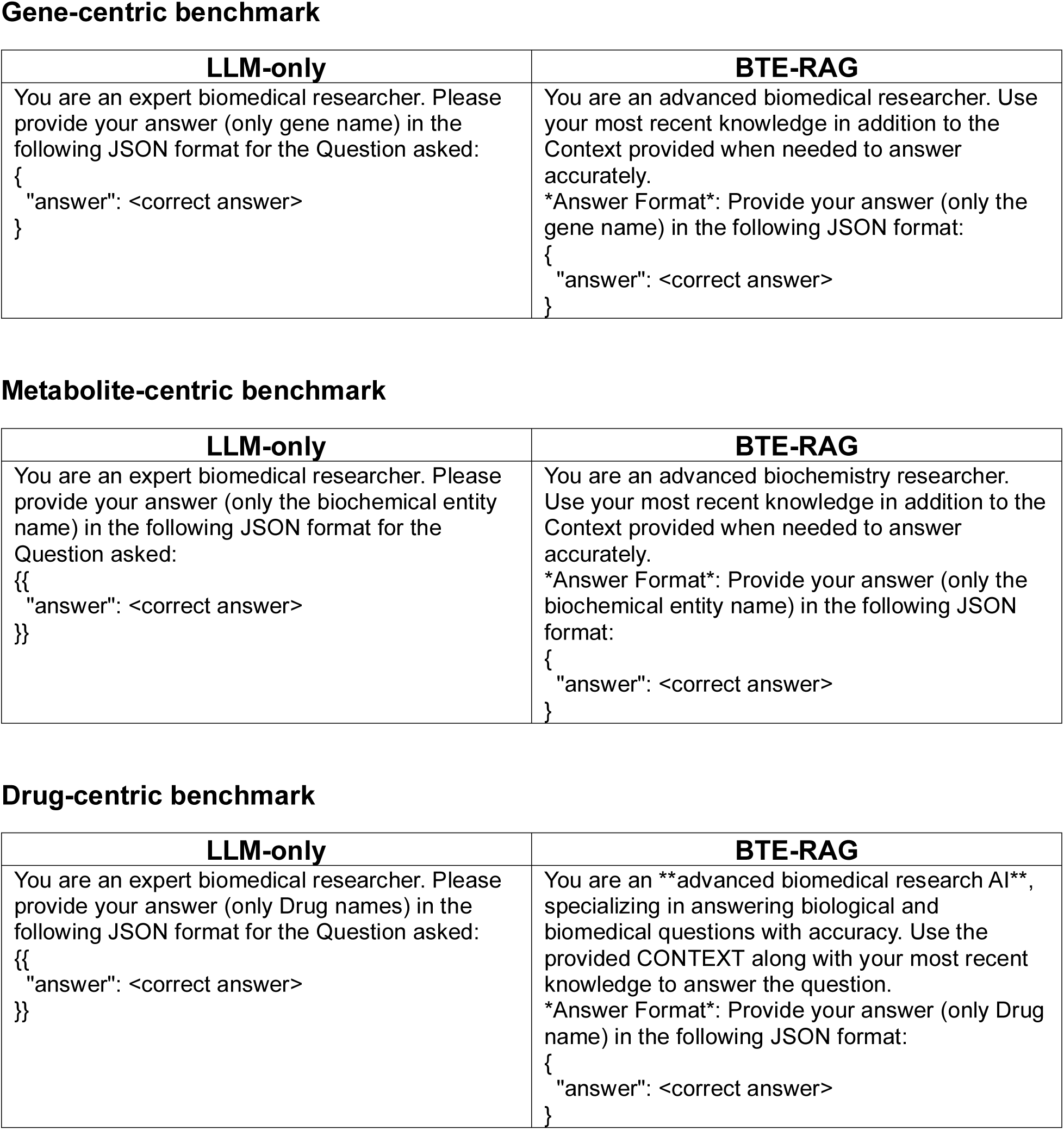
System Prompts Gene-centric benchmark

## References

1. Hou, W. & Ji, Z. Assessing GPT-4 for cell type annotation in single-cell RNA-seq analysis. Nature Methods 2024 21:8 21, 1462–1465 (2024).

2. Rives, A. et al. Biological structure and function emerge from scaling unsupervised learning to 250 million protein sequences. Proc Natl Acad Sci U S A 118, e2016239118 (2021).

3. Lin, Z. et al. Evolutionary-scale prediction of atomic-level protein structure with a language model. Science (1979) 379, 1123–1130 (2023).

4. Meier, J. et al. Language models enable zero-shot prediction of the effects of mutations on protein function. in Proceedings of the 35th International Conference on Neural Information Processing Systems (Curran Associates Inc., Red Hook, NY, USA, 2021).

5. Zheng, Y. et al. Large language models in drug discovery and development: From disease mechanisms to clinical trials. arxiv.orgY Zheng, HY Koh, M Yang, L Li, LT May, GI Webb, S Pan, G ChurcharXiv preprint arXiv:2409.04481, 2024•arxiv.org.

6. Miller, K. et al. Dynamic few-shot prompting for clinical note section classification using lightweight, open-source large language models. Journal of the American Medical Informatics Association 32, 1164–1173 (2025).

7. Ji, Z. et al. Survey of Hallucination in Natural Language Generation. ACM Comput Surv 55, (2023).

8. Vaswani, A., et al. Attention Is All You Need (Transformer Architecture). (2023).

9. Kim, Y. et al. Medical Hallucinations in Foundation Models and Their Impact on Healthcare. (2025).

10. Idnay, B. et al. Environment scan of generative AI infrastructure for clinical and translational science. npj Health Systems 2, 1–11 (2025).

11. Ibrahim, M. et al. Generative AI for synthetic data across multiple medical modalities: A systematic review of recent developments and challenges. Comput Biol Med 189, 109834 (2025).

12. Maynez, J., Narayan, S., Bohnet, B. & McDonald, R. On Faithfulness and Factuality in Abstractive Summarization. Proceedings of the Annual Meeting of the Association for Computational Linguistics 1906–1919 (2020) doi:10.18653/v1/2020.acl-main.173.

13. Yang Bs, Y., Jin, Q., Huang Phd, F. & Lu, Z. Adversarial Attacks on Large Language Models in Medicine. (2024).

14. Luo, R. et al. BioGPT: generative pre-trained transformer for biomedical text generation and mining. Brief Bioinform 23, (2022).

15. Kojima, T., Gu, S. S., Reid, M., Matsuo, Y. & Iwasawa, Y. Large Language Models are Zero-Shot Reasoners. Adv Neural Inf Process Syst 35, (2022).

16. Brown, T. B. et al. Language Models are Few-Shot Learners. Adv Neural Inf Process Syst 2020-**December**, (2020).

17. Lewis, P. et al. Retrieval-Augmented Generation for Knowledge-Intensive NLP Tasks. Adv Neural Inf Process Syst 2020-**December**, (2020).

18. Izacard, G. & Grave, E. Leveraging Passage Retrieval with Generative Models for Open Domain Question Answering. EACL 2021 - 16th Conference of the European Chapter of the Association for Computational Linguistics, Proceedings of the Conference 874–880 (2020) doi:10.18653/v1/2021.eacl-main.74.

19. Karpukhin, V. et al. Dense Passage Retrieval for Open-Domain Question Answering. EMNLP 2020 - 2020 Conference on Empirical Methods in Natural Language Processing, Proceedings of the Conference 6769–6781 (2020) doi:10.18653/v1/2020.emnlp-main.550.

20. Zhang, G. et al. Leveraging long context in retrieval augmented language models for medical question answering. NPJ Digit Med 8, 239 (2025).

21. Soman, K. et al. Biomedical knowledge graph-optimized prompt generation for large language models. Bioinformatics 40, (2024).

22. Hou, W., BioRxiv, Z. J.-& 2023, undefined. GeneTuring tests GPT models in genomics. *biorxiv.orgW Hou*, Z JiBioRxiv, 2023•*biorxiv.org*doi:10.1101/2023.03.11.532238.ABSTRACT.

23. Bizon, C. et al. ROBOKOP KG AND KGB: Integrated Knowledge Graphs from Federated Sources. J Chem Inf Model 59, 4968 (2019).

24. Mungall, C. J. et al. The Monarch Initiative: An integrative data and analytic platform connecting phenotypes to genotypes across species. Nucleic Acids Res 45, D712–D722 (2017).

25. Pan, S. et al. Unifying Large Language Models and Knowledge Graphs: A Roadmap. IEEE Trans Knowl Data Eng 36, 3580–3599 (2024).

26. Evangelista, J. E. et al. Toxicology knowledge graph for structural birth defects. Communications Medicine 2023 3:1 3, 1–14 (2023).

27. Callaghan, J. et al. BioThings Explorer: a query engine for a federated knowledge graph of biomedical APIs. Bioinformatics 39, (2023).

28. Carbon, S. et al. The Gene Ontology resource: enriching a GOld mine. Nucleic Acids Res 49, D325–D334 (2021).

29. Knox, C. et al. DrugBank 6.0: the DrugBank Knowledgebase for 2024. Nucleic Acids Res 52, D1265–D1275 (2024).

30. Fecho, K. et al. Progress toward a universal biomedical data translator. Clin Transl Sci 15, 1838–1847 (2022).

31. Gonzalez-Cavazos, A. C. et al. DrugMechDB: A Curated Database of Drug Mechanisms. Sci Data 10, 1–7 (2023).

32. OpenAI. GPT-4o System Card. (2024).

33. Deka, P., Jurek-Loughrey, A. & Padmanabhan, D. Improved methods to aid unsupervised evidence-based fact checking for online health news. Journal of Data Intelligence 3, 474–505 (2022).

34. Deka, P., Jurek-Loughrey, A. & P, D. Evidence extraction to validate medical claims in fake news detection. SpringerP Deka, A Jurek-Loughrey, DPInternational conference on health information science, 2022•Springer **13705 LNCS**, 3–15 (2022).

35. Unni, D. R. et al. Biolink Model: A universal schema for knowledge graphs in clinical, biomedical, and translational science. Clin Transl Sci 15, 1848 (2022).

36. Wu, C., MacLeod, I. & Su, A. I. BioGPS and MyGene.info: organizing online, gene-centric information. Nucleic Acids Res 41, D561–D565 (2013).

37. OpenAI et al. GPT-4 Technical Report. (2023).

38. Chen, Y. et al. Iterative Prompt Refinement for Mining Gene Relationships from ChatGPT. *bioRxiv* 2023.12.23.573201 (2023) doi:10.1101/2023.12.23.573201.

39. Marvin, G., Hellen, N., Jjingo, D. & Nakatumba-Nabende, J. Prompt Engineering in Large Language Models. 387–402 (2024) doi:10.1007/978-981-99-7962-2_30.

40. Sahoo, P., et al. A Systematic Survey of Prompt Engineering in Large Language Models: Techniques and Applications. (2024).

41. Li, M., Kilicoglu, H., Xu, H. & Zhang, R. BiomedRAG: A retrieval augmented large language model for biomedicine. J Biomed Inform 162, 104769 (2025).

42. Morris, J. H. et al. The scalable precision medicine open knowledge engine (SPOKE): a massive knowledge graph of biomedical information. Bioinformatics 39, (2023).

43. Jin, Q. et al. Pubmedqa: A dataset for biomedical research question answering. arxiv.orgQ Jin, B Dhingra, Z Liu, WW Cohen, X LuarXiv preprint arXiv:1909.06146, 2019•arxiv.org.

44. Kwiatkowski, T. et al. Natural Questions: A Benchmark for Question Answering Research. Trans Assoc Comput Linguist 7, 452–466 (2019).

45. Davis, A. P. et al. Comparative Toxicogenomics Database (CTD): update 2023. Nucleic Acids Res 51, D1257–D1262 (2023).

46. Schriml, L. M. et al. Disease Ontology: a backbone for disease semantic integration. Nucleic Acids Res 40, D940–D946 (2012).

47. Bateman, A. et al. UniProt: the Universal Protein Knowledgebase in 2025. Nucleic Acids Res 53, D609–D617 (2025).

48. Pilarczyk, M. et al. Connecting omics signatures and revealing biological mechanisms with iLINCS. Nat Commun 13, 4678 (2022).

